# A Comprehensive Framework for Spatio-Temporal Analysis of DNA Damage Foci in Tumor Spheroids

**DOI:** 10.1101/2024.07.19.604255

**Authors:** Martine Cazalès, Théo Liu, David Bernard, Denis Jullien, Corinne Lorenzo

## Abstract

The evaluation of DNA damage response, particularly DNA damage foci formation, is crucial for understanding tumor biology and assessing the impacts of various drugs. We have developed a sophisticated semi-automated image analysis pipeline which generates quantitative map of the spatiotemporal distribution of DNA damage foci within live tumor spheroids. Our framework seamlessly integrates live imaging of tumor spheroids via Light Sheet Fluorescence Microscopy with a DNA damage foci formation assay using a genetically encoded fluorescently labeled DNA damage sensor. By combining advanced imaging techniques with computational tools, our framework offers a powerful tool for studying DNA damage response mechanisms in complex 3D cellular environments.

**MOTIVATION:** The motivation of this work is to propose a comprehensive framework that facilitates the study of DNA repair mechanisms within 3D contexts, specifically using tumor spheroid models. By integrating advanced imaging technologies and genetically encoded fluorescent sensors, our goal is to offer researchers a robust methodology for observing and analyzing DNA damage dynamics in realistic tissue-like environments. This framework is designed to enhance accessibility and streamline data processing, thereby empowering the scientific community to investigate DNA repair processes in 3D with greater precision and efficiency.

## Introduction

Genome integrity is constantly threatened by DNA damage triggered by internal and external factors. To counteract these challenges, cells employ a complex network of DNA Damage Response (DDR) pathways, activating DNA repair systems, cell cycle checkpoints, and apoptosis [1]. Dysfunctions in these DDR mechanisms can lead to the accumulation of DNA damage, contributing to a spectrum of diseases including cancer, immune disorders, cardiovascular issues, and neurological degeneration [2].

Advancements in studying DNA repair dynamics within 3D environments have leveraged advanced microscopy techniques and specialized analysis tools. High-resolution methods like 3D-structured illumination microscopy have significantly enhanced spatial resolution, revealing intricate structures of DNA repair foci and the organizational dynamics of repair proteins within them [3]. Complementary multi-scale imaging approaches provide insights across various cellular organizational levels, from individual foci to larger nuclear structures [4]. Live-cell microscopy, particularly leveraging fluorescently tagged proteins such as GFP, enables real-time observation of DNA repair kinetics [5]. This allows for dynamic tracking of repair protein recruitment to damage sites and the temporal evolution of repair foci. Besides, sophisticated image analysis algorithms like FocAn have been developed to automate the detection and quantification of DNA damage markers, including γH2AX foci, in 3D confocal datasets [6], crucial for managing and interpreting large imaging datasets accurately.

Despite these advancements, it’s important to acknowledge that current techniques predominantly focus on the cellular scale, thereby limiting their direct applicability to *in vivo* conditions. Our understanding of DDR mechanisms, especially DNA repair systems, has been primarily derived from observations in 2D cell cultures. However, the translation of these findings to tissue or organ contexts remains relatively unexplored. To bridge this gap, 3D tumor spheroid (TS) models are increasingly employed to study DDR within more physiologically relevant environments [7–10]. These models mimic the structural complexity of solid tumors, with distinct layers such as proliferating, quiescent, and necrotic regions [11]. These layers give rise to gradients in essential factors such as oxygen, nutrients, and pH levels, providing a more realistic simulation of the tumor microenvironment.

Advanced imaging technologies like Selective Plane Illumination Microscopy (SPIM), a variant of Light Sheet Fluorescence Microscopy, facilitate high-resolution, long-term live imaging of cellular processes within intact 3D cell culture models while minimizing phototoxicity [12–15].

Our proposed framework extends these methodologies by dynamically analyzing DNA damage foci formation within TS, aiming to integrate cellular-scale observations with tissue-level understanding of DNA repair processes. By utilizing genetically encoded fluorescent DNA damage sensors combined with SPIM, we monitored changes in nuclei and DNA damage foci concurrently, capturing cellular responses to genotoxic stress over extended periods. This framework integrates a strategic combination of elements, including the DNA damage sensor, microscopy techniques, and advanced computational tools. Central to our approach is a dedicated pipeline that incorporates an original “3D texture” data transformation program and an analytical notebook, designed to streamline data processing and analysis. The pipeline encompasses essential preprocessing steps, precise segmentation, normalization, and the creation of 3D texture maps. Additionally, we have developed a dedicated notebook to facilitate in-depth analysis of these multiparametric datasets, focusing on spatial dimensions, time, and foci/nuclei range. This pipeline is designed to be accessible to individuals with basic knowledge of image processing and analysis and exclusively employs open-source tools. This framework represents a significant advancement in investigating DNA repair mechanisms within complex 3D cell culture models, essential for advancing therapeutic strategies against DNA damage-related diseases.

## Results

### Visualizing DNA Damage Induction Following Chemotherapeutic Treatment in Engineered Tumor Spheroids Using a Fluorescent Sensor

Tumor spheroids (TS) engineered to express a genetically encoded fluorescent DNA damage sensor, specifically the minimal domain (mDomain) of the 53BP1 protein fused with mCherry, were developed. The 53BP1 protein plays a crucial role in recognizing double-strand DNA damage, with its mDomain forming observable nuclear foci under fluorescence microscopy (**Fig. 1** and [16]). In untreated TS, mCherry-mDomain exhibited diffuse nuclear staining and concentrated at discrete nuclear bodies [17] and DNA damage foci (**Fig. 1A**). To induce DNA damage, engineered TS were treated with the chemotherapeutic agent bleomycin, known for inducing both single- and double-strand breaks in DNA [18]. To assess the drug’s impact throughout the TS, we evaluated the foci index —representing the percentage of positive nuclei with more than three foci—at various depths in cleared TS under both untreated and treated conditions. As depicted in **Figure 1B**, an average baseline foci index of 29% was observed in untreated TS, indicating the inherent level of damage. Following bleomycin treatment, a consistent twofold increase in the foci index was observed across all depths of the TS. Thus, the foci index consistently elevated in response to bleomycin treatment, with no significant variation noted in the TS depth. This increase in the foci index after bleomycin treatment demonstrates the drug’s impact on DDR processes throughout the entire TS.

**Figure 1:**
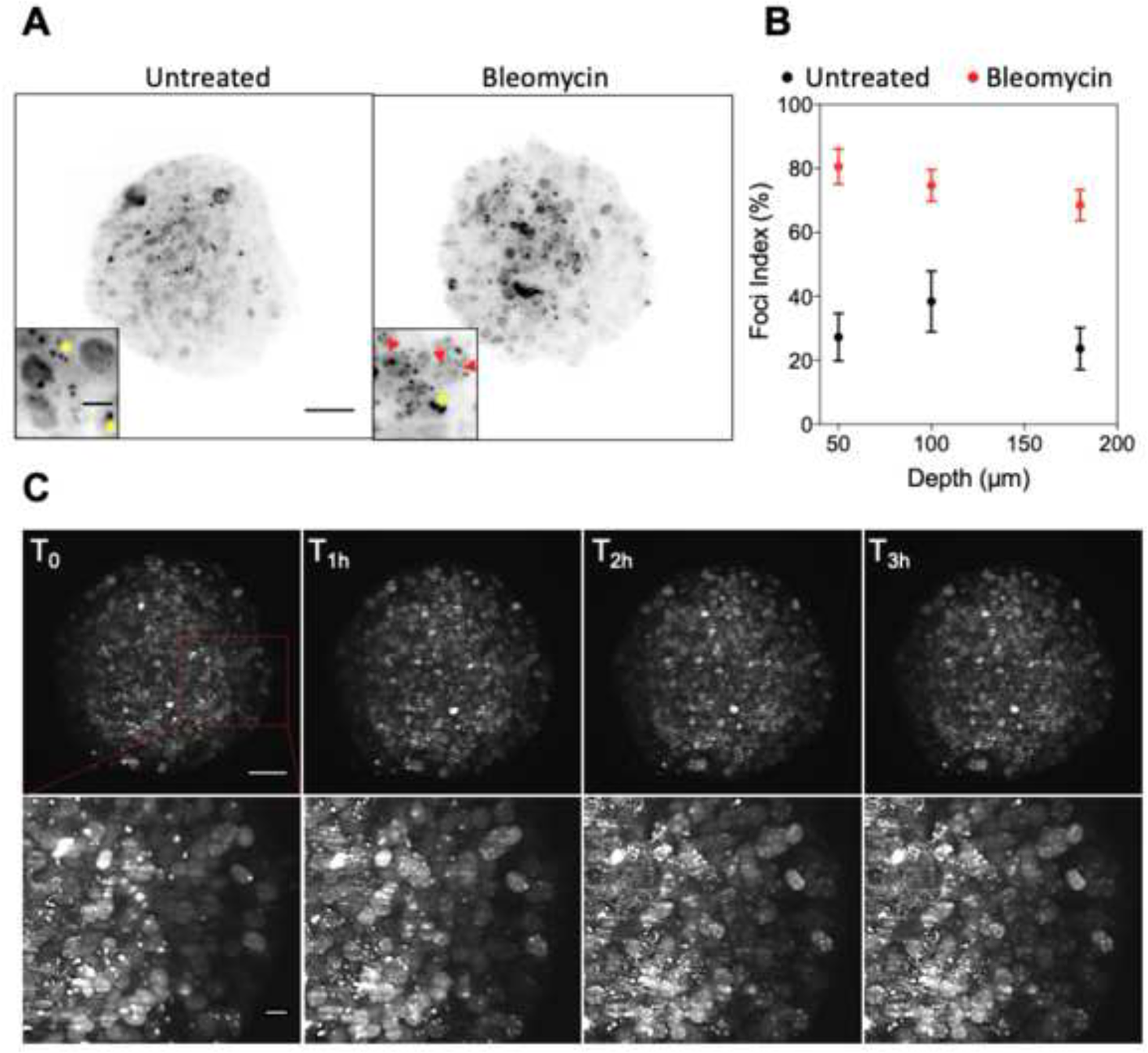
Visualization of DNA Damage Foci in Tumor Spheroids Using Fluorescent DNA Damage Sensor mCherry-mDomain. **(A)** Representative Light Sheet Microscopy images of cleared TS expressing the mCherry-labeled 53BP1 minimal DNA damage sensor domain (mDomain), which exhibits diffuse nuclear staining and concentrates at discrete nuclear bodies (yellow asterixis) and DNA damage foci (red arrows) in untreated and bleomycin conditions. Scale bar 100 µm and inset 10 µm **(B)** DNA damage foci index, representing the percentage of nuclei with more than three foci, is quantified in untreated and bleomycin-treated TS at different depths. **(C)** Short-term live-cell imaging of TS expressing the mCherry-mDomain after bleomycin treatment. The bottom arrow displays a zoomed-in region highlighted in the top square red. Time is indicated in hours, and the scale bar represents 100 µm (top) 10 μm (bottom).

### Optimizing Live Light Sheet Imaging for Long-Term Observation of DNA Damage in Engineered Tumor Spheroids

3D time-series images of engineered TS treated with or without bleomycin were captured using a custom-built SPIM, as previously described by Lorenzo *et al.,* [12]. The TS were cultured in a chamber filled with physiological medium within an agarose container to maintain long-term viability [19]; The mCherry-mDomain protein was excited with a 595 nm laser, and Leica 10x/0.3 and 20x/0.5 water immersion objectives were used for excitation and emission, respectively. Emitted light was collected through a 594-nm long-pass detection filter and a Hamamatsu Orca Flash 4.0 sCMOS camera. Imaging parameters, including laser power, exposure time, and interval time, were optimized to achieve a balance between signal-to-noise ratio, imaging power, and phototoxicity, thereby providing 3D datasets with sufficient temporal and spatial resolution to monitor DNA damage foci formation induced by bleomycin over 2-3 days. Indeed, optimizing live imaging over extended periods while minimizing unwanted phototoxic effects, in particular DNA damage, was crucial for investigating DDR to obtain more accurate and relevant observations of DNA damage foci formation induced by the bleomycin treatment. One hour after bleomycin introduction, we observed a gradual increase in spot appearance over time, corresponding to DNA damage induced by bleomycin treatment and the recruitment of mDomain to the DNA break sites (**Fig. 1C and Video S1**). Conversely, no such increase was observed in untreated TS (**data not shown**).

### Standardized Pipeline for Streamlined 4D DNA Damage Mapping and multiparametric analysis

Creating a quantitative 4D map of DNA damage foci within TS presents several complex challenges, particularly in the context of long-term imaging experiments. For instance, capturing 150 z stacks every hour over a span of 50 hours results in a substantial data volume of 125 GB per experiment. Furthermore, conducting up to 10 individual experiments per condition can contribute to a significant data load of 1 TB. Managing multiple terabytes of data becomes even more intricate when dealing with diverse experimental conditions, including controls and various drug treatments.

The task of extracting a 4D quantitative map of DNA damage involved measuring the number of foci/nuclei in both spatial and temporal dimensions. A significant challenge encountered was identifying nuclei and spots with a unique label, herein the mCherry-mDomain, within dense TS. Nuclei, characterized by low intensity and large size, contrasted with DNA damage foci, which exhibited high intensity and small size (**Fig. 1 A, C and Video S1**). Despite the challenges posed by segmenting nuclei with low intensity levels and poorly defined contours, we prioritized minimizing phototoxicy. This decision was crucial to prevent DNA damage induced by multiple illuminations, while also aiming for accurate segmentation of nuclei. In contrast, DNA damage foci proved more amenable to segmentation due to their high intensity fluorescence and discrete structure.

We implemented a comprehensive pipeline designed to efficiently handle large volumes of data. This approach encompassed preprocessing, object identification, data transformation, visualization, and analysis (**Fig.2 and 3**).

**Figure 2:**
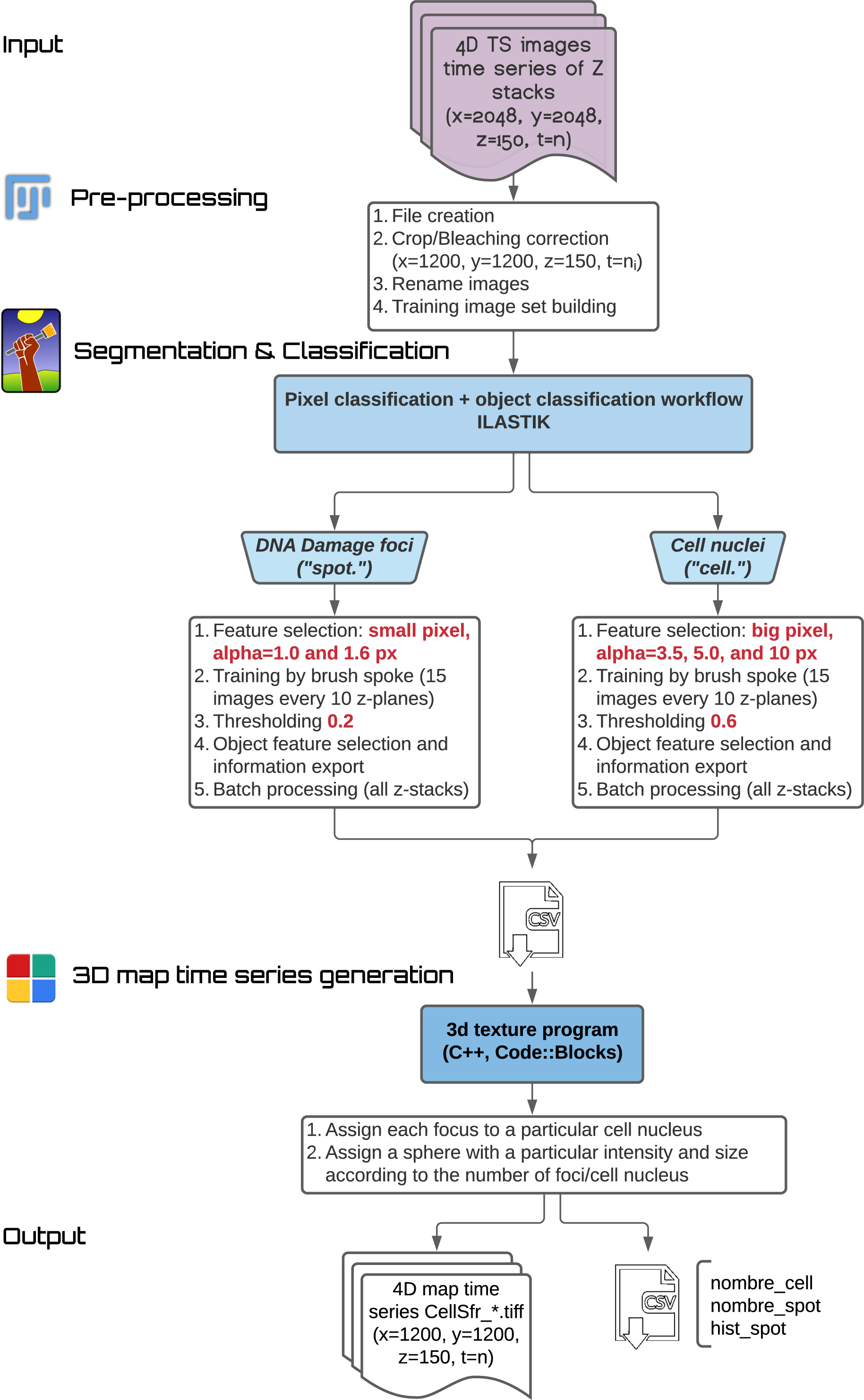
Workflow for Generating 3D Time Series Maps of DNA Damage Foci in Tumor Spheroids. **(A)** 4D TS images are captured over time in Z-stacks, cropped and processed for bleaching correction. **(B)** ILASTIK is utilized for pixel and object classification to identify DNA damage foci (“spots”) and cell nuclei (“cells”). **(C)** A custom 3D texture program assigns each focus to a specific cell nucleus. **(D)** The final output includes quantitative data on cell and spot numbers, along with a histogram of spots per nucleus, resulting in 4D map time series files.

**Figure 3:**
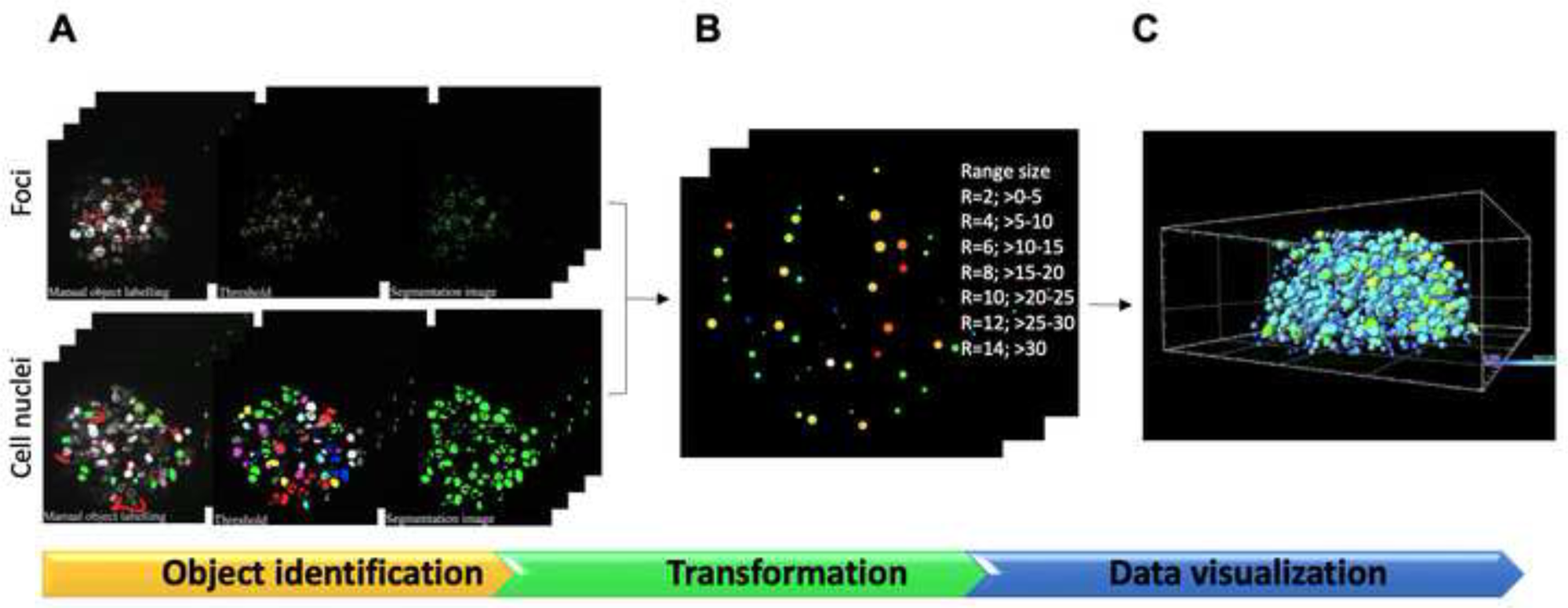
Framework for Identifying and Visualizing DNA Damage Foci in Tumor Spheroids. This concise framework delineates the process for identifying and visualizing DNA damage foci in TS. **(A)** Object identification through preprocessing, pixel classification, and segmentation. **(B)** data transformation utilizing a custom 3D texture program. it defines range sizes for foci visualization based on their number per nucleus. **(C)** Visualization is achieved through 3D map time series, providing insights into the spatial and temporal distribution of DNA damage. it defines range sizes for foci visualization based on their abundance per nucleus.

#### Automated Pre-Processing with a Dedicated FIJI Macro

A dedicated FIJI [20] Macro [crop_bleachcorrect.ijm] was developed to load all 3D time-series images, crop unnecessary parts, and correct bleaching using the simple ratio method [21]. Following this preprocessing step, a training image set was created to identify foci and cell nuclei. Contiguous z planes (z=75) were extracted from two representative pre-processed 3D time-point images—one with bleomycin-induced DNA damage foci and one without. To streamline the computational process, both z stacks were concatenated and cropped again to reduce calculation time during the training step.

#### Ilastik-Based Pixel and Object Classification for Batch Processing: Training, Annotation, and Application to 3D Time-Series Images

Utilizing Ilastik [22], a Random Forest classifier was trained with selected features specific to each object of interest (*e.g*., small pixels for foci and large pixels for cell nuclei, **Fig. 2**). Image pixels were categorized into two classes—background and object of interest—manually labeled using the paintbrush tool (**Fig. 3A**). Approximately 15 images distributed throughout the training set (every 10 z planes) were fully annotated. New pixel classifications were added until stable probability maps that adequately distinguished foci or cell nuclei were obtained. Subsequently, a threshold was applied to segment and extract various features of each identified object (**Fig. 2**), including centers, radii, bounding boxes, and mean intensity.

The trained classifier was then applied to all pre-processed 3D time-series images as a batch processing operation, taking approximately 8 hours for a 3D image time series of 50 points. The results of object classification were saved in the working directory as CSV tables and hdf5 files. Despite challenges such as irregular shapes, poorly defined boundaries, low and inhomogeneous contrast, and considerable noise in the images of cell nuclei, the Ilastik workflow demonstrated its capability to yield acceptable segmentations for subsequent 3D time-series map generation, as illustrated in **Figure 3A**.

#### Foci and Nuclei Localization Coupling in 4D Time-Series Map Generation with the ‘3D Texture’ Program

The “3d texture” program encompasses four key global functions: “genere_rayon_fix_cell,” “genere_SFR,” “genere_BBOX,” and “lire_csv.” Initially, the program redefined objects based on their stored features, including centers, radii, bmin, bmax, and intensity. These objects were classified as either “cell” (representing cell nuclei) or “spot” (indicating DNA damage foci) based on specific conditions:

~~~
If ((bDim0.320.32<6000 && bDim0.320.32>100)) /// cell
If ((bDim0.320.32<100 && bDim0.320.32>0)) /// spot
~~~

Here, bDim represents the dimensions of the bounding box in both the x and y directions, and 0.32 denotes the x and y pixel size (in µm). Subsequently, the program conducts a test to determine if a spot belongs to a cell, utilizing an object-object intersection test in a 3D space. The axis-aligned bounding boxes (AABB) technique is employed for this test, comparing the spot and cell bounding boxes in three dimensions (x, y, and z axes). A spot is considered contained within a cell if:

~~~
Xmin_spot > Xmin_cell ∧ Ymin_spot > Ymin_cell ∧ Zmin_spot > Zmin_cell ∧ Xmax_spot < Xmax_cell
∧ Ymax_spot < Ymax_cell ∧ Zmax_spot < Zmax_cell
~~~

The program then calculates the number of spots belonging to a cell, assigning a 3D sphere of radius RvRv based on the range of spots contained in the cell (**Fig. 3B**). Additionally, the program computes the center and radius of the TS by determining the minimum and maximum positions of the cell centers in all three dimensions. Euclidean distances from each cell to the TS center are calculated to position each attributed 3D sphere with radius RvRv in this new 3D coordinate system (**Fig. 3B**).

The output files include:

- The 3D time-series map (“CellSfr_*.tif”)
- Two CSV files named “nombre_cell.csv” and “nombre_spot.csv,” reporting the number of cell nuclei and DNA damage foci over time
- A “hist_spot” file providing the number of DNA damage foci as a function of TS layer and time. Each layer represents a distance of 0.1RvRv (0.1R) from the spheroid center.

During post-processing, a FIJI macro named “fusion_volumes3D” combines the 4D maps generated from multiple experiments, creating a unified dataset that consolidates data from up to four experiments into a single virtual experiment (**Fig.3C**). Following this fusion process, the merged dataset can be seamlessly imported into 3D visualization and analysis software like IMARIS, as depicted in **Figure 3C** and **Video S2, S3**, and **S4**. This transformation reduces the dataset size from 125GB to 22GB, greatly enhancing its manageability for analysis across platforms, including FIJI and others.

#### In-Depth Multiple-Parametric Analysis of DNA-Damaged Cells

For an in-depth multiple-parametric analysis of DNA-damaged cells/nuclei, a FIJI macro “3D measurement” and a Jupyter notebook “nuclei/foci analysis” are provided to analyze measurement data from multiple CSV files extracted from the transformed dataset (**Fig. 4**). The macro processes each transformed 3D time-series tiff file in an input directory by opening the file and applying a 3D watershed algorithm for image segmentation (**Fig. 4A**). It uses the 3D Manager [23] to handle and measure the segmented 3D images, saving the measurement results as CSV files in an output directory. The script then closes the results window and repeats the process for all TIFF files, enabling batch processing for 3D time-series transformed image.

**Figure 4:**
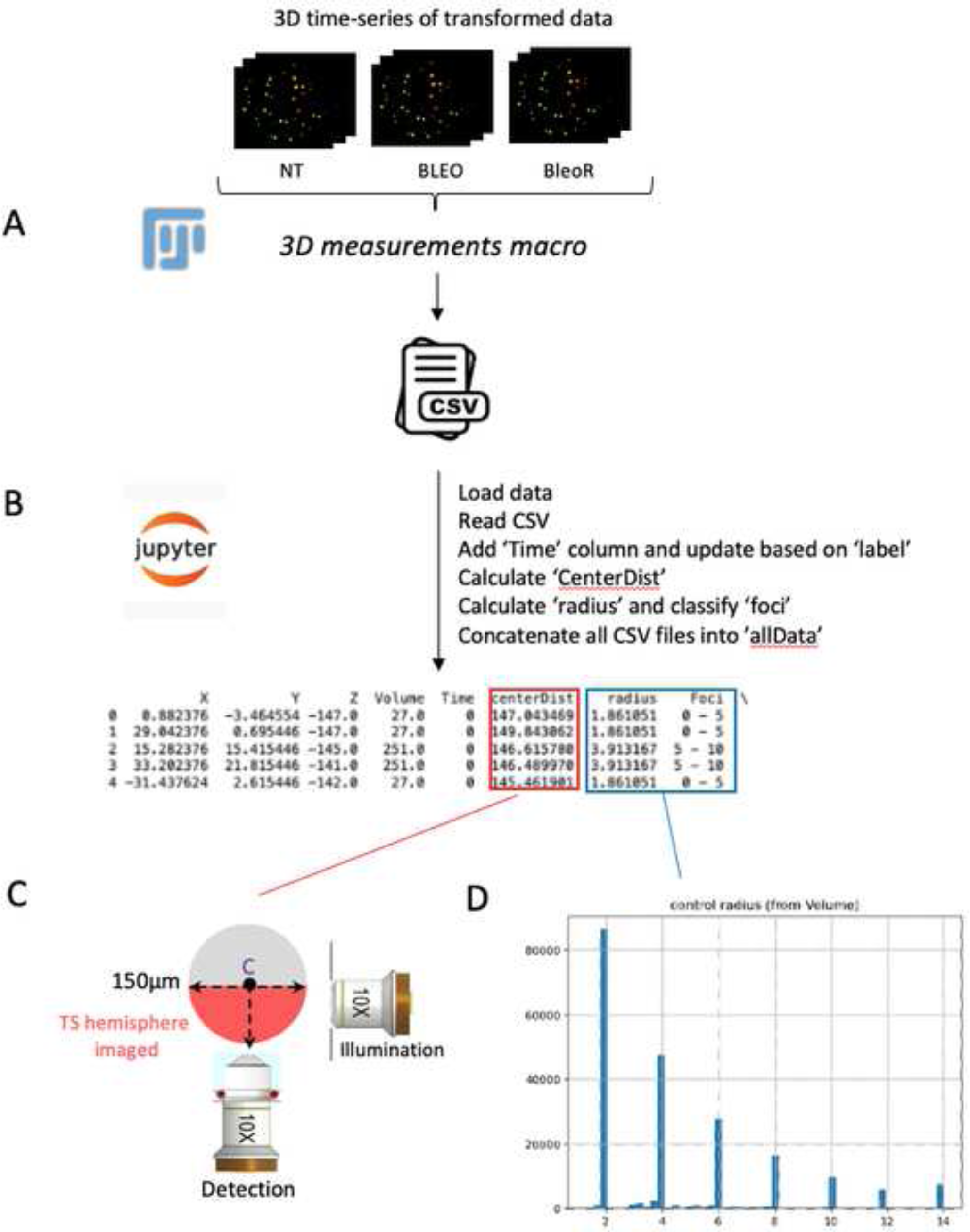
Data Transformation and Analysis Pipeline. **(A)** 3D Measurements Macro: FIJI macro segments and measures 3D TIFF files, saving results as CSV. **(B)** Data Processing: Jupyter notebook preprocesses CSV data, organizes by time, computes distances and radii. **(C)** Schema: Filters data by spatial range (centerDist < 150). **(D)** Graph Example: Classifies foci/nuclei by size intervals, normalizes with distGroup for spatial analysis.

The Jupyter notebook includes a function called *loadData*, which facilitates the loading and preprocessing of each CSV file. This preprocessing involves re-organizing data by time, adjusting spatial coordinates, computing distances and radii, and categorizing the data accordingly.

After preprocessing, the data is consolidated into a single DataFrame named *allData* (**Fig. 4B**). The calculation of *centerDist* using the Euclidean distance formula and subsequent filtering ensured that only data points within a relevant spatial range (*centerDist* < 150, **Fig. 4C**) were retained for further analysis. The derivation of radius for each data point allowed for the classification of foci/nuclei into intervals (foci) based on their size, providing insights into the distribution of DNA damage foci (**Fig. 4D**).

To enhance data interpretability, a new column named *distGroup* is introduced. This column groups data points corresponding to foci/nuclei into distance bins based on quantiles, with distinct labels. Normalization ensures that each distance group contains a comparable number of cells/nuclei, mitigating biases arising from the hemispherical shape of the data distribution. Dividing the data into equally-sized groups based on distance from the center allows for a more accurate representation of cell/nuclei distribution and behavior over time and under different conditions. This stratification facilitates meaningful comparisons across conditions, as each distance group represents a consistent portion of the cell population.

Subsequently, a categorical plot is generated to visualize the count of foci/nuclei within each distance group, stratified by different experimental conditions. Additionally, the notebook generates a series of heatmaps arranged in a grid format, illustrating the mean count of foci/nuclei over time for various distance groups and foci ranges across different conditions. This visual representation aids in understanding the distribution of foci/nuclei across spatial and temporal dimensions, revealing patterns and disparities among the experimental conditions.

### Showcasing DNA Damage Response Dynamics in Tumor Spheroids

To demonstrate our proof of concept, we employed two distinct bleomycin treatments. The first involved continuous exposure of TS to bleomycin at a concentration of 10 µg/ml (referred to as BLEO). The second method entailed exposing TS to bleomycin for two and a half hours (referred to as BleoR), also at 10 µg/ml, followed by thorough washing and imaging in a physiological chamber filled with culture medium.

Comprehensive kinetic and statistical analyses were conducted using our 3D texture and Jupyter notebook analysis programs to delve deeply into DNA damage foci within TS, as illustrated in Figures 5 and 6. Initially, we presented the count of cells/nuclei (**Fig. 5A**) and spots (**Fig. 5B**) across the entire TS over time, along with the average number of foci per nucleus (**Fig. 5C**). No significant difference in the count of identified cells was observed among untreated TS, BLEO, and BleoR treated TS. However, as expected, both BLEO and BleoR treatments significantly increased the number of detected spots and the average number of foci per nucleus, particularly noticeable between 3-5 hours post-treatment initiation (P-value < 0.05 **, **Fig. 5B, C**). A plateau was reached, showing a peak of approximately 15 ± 5 average foci per nucleus, followed by a gradual decline towards basal levels observed in the control condition (**Fig. 5C**).

**Figure 5:**
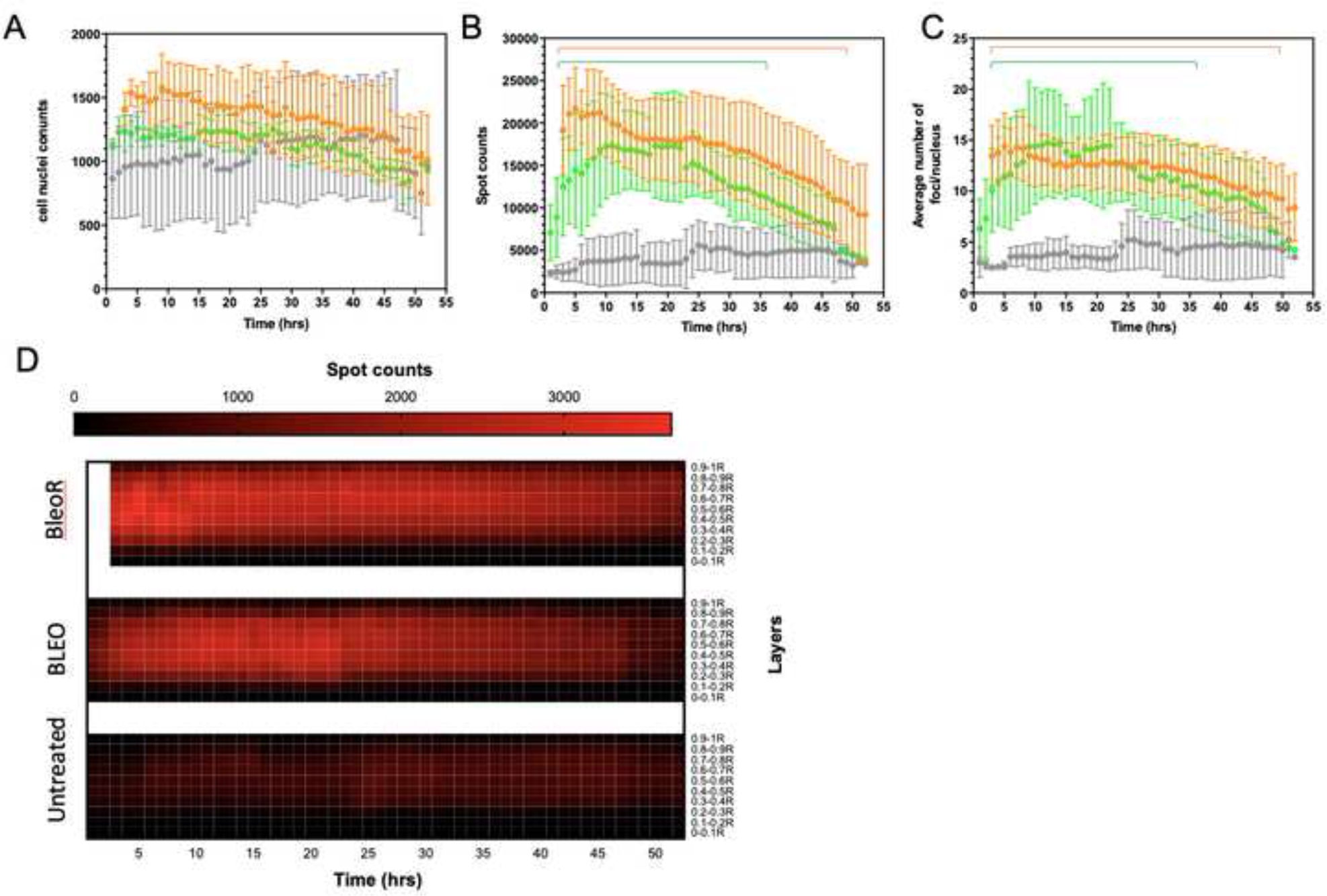
Overall analysis of Foci Distribution Over Time and Throughout the TS Under Different Experimental Conditions. Graphs plot the total number of cells/nuclei **(A)**, total number of spots/foci **(B)**, and average number of foci per nucleus **(C)** over time. Mean curves are estimated from n=4 independent experiments. Error bars represent standard deviation. Statistical significance was determined using multiple unpaired t-tests, with P< 0.05 indicating significance (comparing BleoR and BLEO vs. control) representing by a line. Heatmaps depict the density of spots over time and distance from the TS center under different experimental conditions.

**Figure 6:**
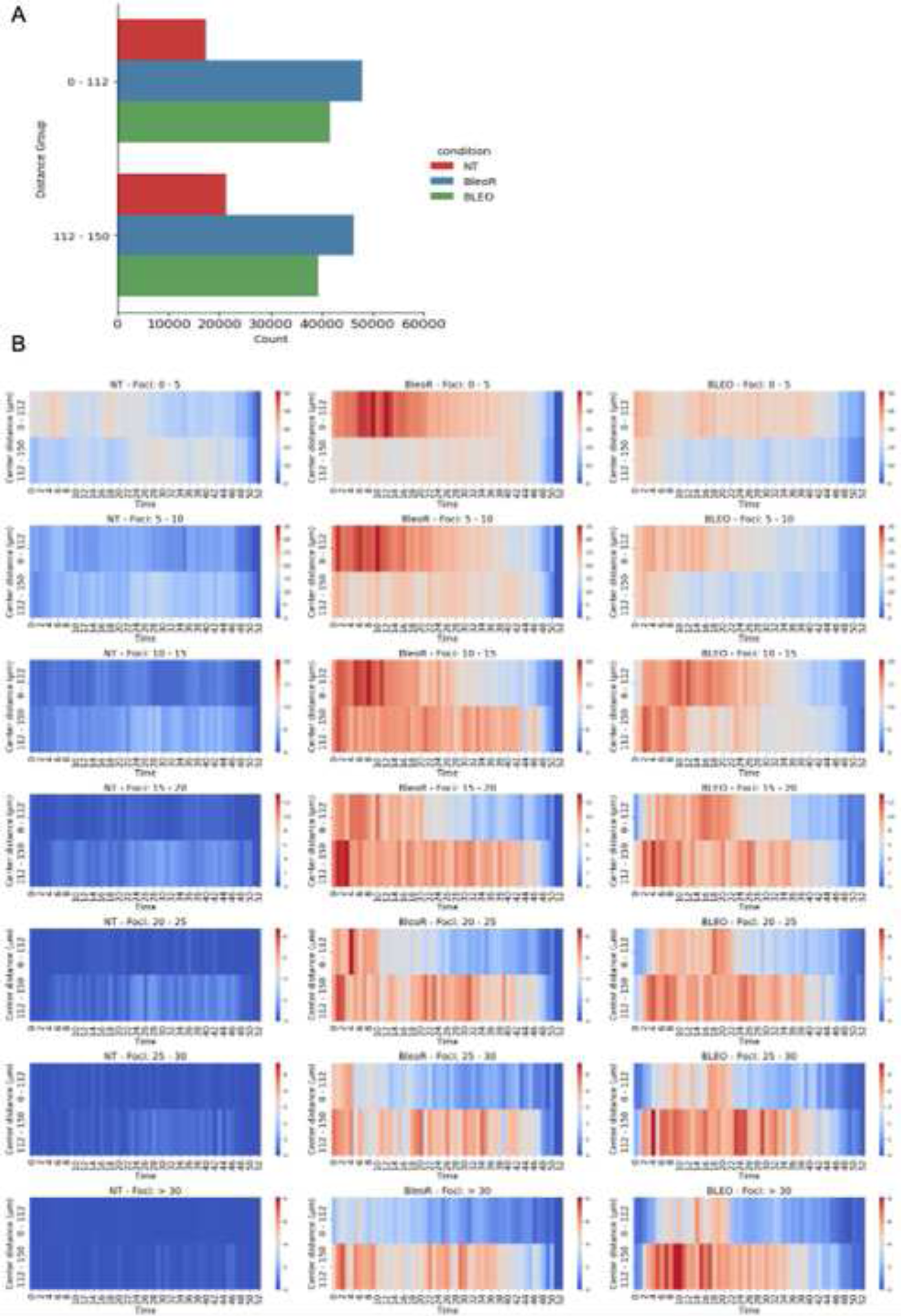
Spatiotemporal Distribution of DNA-Damaged Cells/Nuclei Within Foci Ranges Under Different Treatment Conditions. **(A)** Graph categorizes foci/nuclei within TS into two layers based on their distance from the center: an inner layer (0-112 µm) representing those closer to the center and an outer layer (112-150 µm) for those further away. This categorization utilizes the distGroup column for accurate spatial stratification. **(B)** Heatmaps illustrate the spatiotemporal distribution of DNA-damaged cells/nuclei over time across different foci ranges and treatment conditions. Rows depict specific foci ranges, while columns denote various treatment conditions. Color intensity indicates the number of damaged cells/nuclei, with darker colors signifying higher counts.

Furthermore, following both treatments, the number of DNA damage foci per cell was significantly higher compared to untreated cells, where an average of 2.3 ± 0.15 foci per nucleus was observed within the investigated time frame (**Fig. 5C**). This observation underscores the effectiveness of bleomycin treatments in inducing DNA damage foci formation, leading to a distinct and sustained response compared to the untreated control.

The kinetics of DNA damage foci formation did not exhibit a significant difference between the two types of bleomycin treatments. However, a subtle distinction emerged in their behavior. Specifically, under BLEO conditions, the decay of the foci number was more pronounced, returning closer to basal levels within 37 hours after treatment initiation. Conversely, with BleoR conditions, the decay was less evident, maintaining a significant difference from the control basal level even at the 50-hour mark. This nuanced difference becomes more apparent when comparing the heatmaps (**Fig. 5D**), which depict the dynamic changes in spot or foci numbers over time, stratified across distinct layers of TS. The color intensity in each cell of the heatmap represents the quantity of spots observed at the intersection of a specific time point and TS layer.

Upon careful examination, notable patterns emerge, indicating variations in spot distribution across both temporal and spatial dimensions. Peaks in color intensity at specific time points suggest rapid responses, with stabilization over time. Certain layers consistently exhibit higher or lower spot counts, suggesting layer-specific responses to treatment.

This nuanced depiction provides valuable insights into the spatiotemporal dynamics of spot formation within different layers of TS, highlighting the complexity of cellular responses and offering a comprehensive view of treatment impacts across the entire three-dimensional structure of TS. Visualization of virtual experiments for each condition aligns with these findings, showing an initial burst of DNA damage foci for BLEO and BleoR conditions, particularly prominent in outer layers, followed by stabilization and eventual decrease (**Video S2, S3, and S4**).

To deepen our analysis, we divided TS into two layers based on distance from the center, corresponding to proliferative and quiescent layers (**Fig. 6A, B**). Heatmaps in **Figure 6C** illustrate foci/nuclei density over time and distance from the TS center across different experimental conditions (NT, BleoR, BLEO), categorized by distinct foci ranges per nucleus. Each row represents specific foci ranges, while columns correspond to the three conditions. The x-axis indicates time (0 to 52 minutes), and the y-axis represents distance from the center, with color intensity indicating the number of damaged cells/nuclei.

Across all conditions, damaged cells/nuclei generally increase over time, particularly at greater distances from the center corresponding to TS outer or proliferative layers. NT shows minimal initial damage, with gradual increase, especially at higher distances and lower foci ranges. BleoR displays a significant increase in damaged cells/nuclei over time, especially in the 0-5 and 5-10 foci ranges, with notable activity in higher ranges as well. BLEO exhibits the highest damage levels, particularly in lower foci ranges, with a strong increase over time and highest counts in outer layers. Both BleoR and BLEO significantly increase foci formation compared to NT, with BLEO showing a more potent, dose-dependent effect.

Treated conditions demonstrate higher and more variable densities of damaged cells/nuclei, with BleoR showing high density and variability, particularly in outer layers of TS, while BLEO demonstrates high but more uniform density, suggesting subtle differences in response to continuous versus transient exposure. These findings underscore the substantial impact of bleomycin on DNA damage, with pronounced effects under continuous treatment, highlighting cellular response dynamics to the drug.

## Discussion

This work presents an innovative framework for investigating DNA damage within TS, addressing a critical challenge in cancer research: monitoring DDR in a 3D context. Unlike traditional 2D cell cultures, which offer valuable insights into DDR mechanisms but lack the complexity of real tumor environments, TS provide a more physiologically relevant model by mimicking solid tumor properties. The study’s framework combines SPIM with a genetically encoded fluorescent DNA damage sensor to track DNA damage foci within live TS. This integration of advanced imaging techniques with a robust pipeline enables the generation of quantitative 4D maps of DNA damage, facilitating detailed spatio-temporal analysis. A significant advancement is the implementation of a semi-automated image analysis workflow, adept at handling the substantial data volumes from prolonged imaging experiments. This pipeline encompasses image pre-processing, segmentation, classification and data transformation steps, providing a comprehensive solution for analyzing and visualizing dynamic cellular events in 4D.

Notably, this framework’s originality lies in its capacity to automate and streamline complex analysis tasks while yielding detailed spatial and temporal insights into DNA damage foci dynamics. Through the use of open-source tools and the provision of a detailed tutorial on GitHub, the pipeline ensures accessibility and reproducibility for other researchers.

**Visualization and Quantification of DNA Damage:** The engineered TS expressing a fluorescent DNA damage sensor provides clear visualization of DNA damage foci. In untreated TS, a baseline foci index is established, while treatment with the chemotherapeutic agent bleomycin results in a consistent twofold increase in the foci index across all TS depths. This highlights the method’s sensitivity and reliability in detecting DNA damage.

**Spatio-Temporal Dynamics:** The framework ability to capture 3D time-series images allows for the observation of DNA damage dynamics over 50 hours. Our analysis reveals that bleomycin treatment induces significant DNA damage uniformly throughout the TS. Moreover, differences in the decay patterns of DNA damage foci are observed between continuous and transient bleomycin treatments, providing insights into the kinetics of DDR processes.

**Layer-Specific Analysis:** The study’s approach also enables the analysis of DNA damage distribution across different TS layers, representing proliferative and quiescent zones. This layer-specific analysis is crucial for understanding how different cellular microenvironments within a tumor respond to DNA damage.

**Data Integration and Analysis:** The creation of 4D maps and the use of a dedicated Jupyter notebook for in-depth data analysis allows for a comprehensive examination of DNA damage patterns. This integration of multiple parameters, such as time, spatial dimensions, and foci count, facilitates a nuanced understanding of the cellular responses to DNA damage.

The developed framework represents a significant step forward in the study of DDR in 3D cell cultures. By providing a more accurate and detailed picture of DNA damage dynamics, this approach can enhance our understanding of how cancer cells respond to genotoxic stress. This is particularly relevant for the development of more effective cancer therapies, as it allows for the evaluation of drug efficacy in a context that closely mimics actual tumors. Future research could expand this framework to study other types of DNA damage and repair processes, as well as to investigate the effects of different chemotherapeutic agents. Additionally, integrating this pipeline with other advanced imaging and computational techniques could further improve the resolution and accuracy of DDR studies in 3D models. In conclusion, the comprehensive framework provides a powerful tool for the spatio-temporal analysis of DNA damage in TS.

## Limitations of the Study

While this study presents significant advancements, it also has several limitations:

1. **Phototoxicity Concerns:** Despite efforts to minimize induced photo-DNA damage, long-term live imaging still introduces phototoxic effects, potentially affecting the accuracy of the results.
2. **Specificity of the DNA Damage Sensor:** The genetically encoded fluorescent DNA damage sensor, while effective, is specific to the 53BP1 protein domain, which may not capture all types of DNA damage responses, potentially overlooking other important repair pathways.
3. **Model Limitations:** Although TS provide a more realistic model than 2D cultures, they still do not fully replicate the complexity of *in vivo* tumor environments, such as the presence of stromal and immune cells, vascularization, and the extracellular matrix.
4. **Temporal Resolution:** The time intervals (1 hour) between image captures, although sufficient for observing general trends, may miss more rapid, transient events in the DNA damage response that occur at shorter timescales.

These limitations highlight areas for potential improvement in future research, such as enhancing imaging techniques to further reduce phototoxicity, developing more sophisticated multi-labeling strategies, and integrating additional cell types and components to better mimic the *in vivo* tumor microenvironment.

## Acknowledgements

This work was supported by the Fondation de la Recherche Médicale (FRM, grant ING20160435179) and by the Conseil de Radioprotection EDF (grant RB 2015-05). We gratefully acknowledge the scientific and technical support of our team members. We also thank L. Casteilla, L. Vaysse and P. Monsarrat for their helpful comments on the manuscript.

## Author Contributions

DJ constructed the mCherry-mDomain and produced the stable transfected HCT116 cell line expressing mCherry-mDomain, which was used for generating tumor spheroids. MC performed the experiments, conducted the light-sheet microscopy acquisition and image analysis. TL designed and wrote the 3D texture code, tested it, and assisted with the development of best-practices workflows and analysis. DB designed and wrote the Jupyter notebook. CL conceived the original idea, provided systematic advice throughout the project, performed the analyses and wrote the manuscript with support from all authors. All authors contributed to the article and approved the submitted version.

## Declaration of Interest

The authors declare no competing interests.

## Declaration of generative AI and AI-assisted technologies in the writing process

During the preparation of this work, the authors used ChatGPT in order to improve the grammatical structure in some paragraphs; After using ChatGPT, the authors reviewed and edited the content as needed and take full responsibility for the content of the publication.

## Supplemental information

**Video S1:** Short-Term Live Imaging of HCT116-mDomain/mCherry Treated with Bleomycin related to Figure 1

**Video S2:** 4D Mapping of DNA Damage in Untreated Tumor Spheroids related to Figures 5 and 6

**Video S3:** 4D Mapping of DNA Damage in Bleomycin-Treated Tumor Spheroids related to Figures 5 and 6

**Video S4:** 4D Mapping of DNA Damage in Bleomycin-Released Tumor Spheroids related to Figures 5 and 6

## STAR Methods

### Cell Culture

HCT116 mCherry-mDomain cells were cultured in DMEM supplemented with GlutaMAX (Gibco), 10% fetal bovine serum (FBS), and 1% penicillin-streptomycin (Pen Strep; Gibco). The cells were maintained in a standard cell culture incubator at 37°C with 5% CO2.

### TS Production

Tumor spheroids (TS) were generated using the centrifugation method detailed by Andilla et al., resulting in uniformly sized spherical TS with a coefficient of variation <10%. For preparation, 96-well plates were pre-coated with 20 mg/ml polyHEMA (Sigma). Cells were seeded at a density of 1500 cells per well in 100 µl of cell culture medium and centrifuged to facilitate TS formation. Following 4–5 days of growth under standard conditions, TS ranging in diameter from 300 to 350 µm were harvested for further experimentation.

### TS Clearing

To achieve TS clearing, 2,2-Thiodiethanol (TDE) was selected as the clearing agent. The protocol involved sequential immersion in baths with increasing concentrations of TDE: 10%, 20%, 30%, 40%, and 50%. Each incubation step was conducted at room temperature with gentle agitation. Specifically, the TS were incubated for 10 minutes in 10% TDE, followed by overnight incubation in 20% TDE. Subsequently, they were incubated for 30 minutes in 30% TDE, 1 hour in 40% TDE, and finally 1 hour in 50% TDE.

### Bleomycin Treatments

TS were incubated with 10 µg/ml of Bleomycin for 2 hours and 30 minutes, followed by extensive washing for 20 minutes before being placed into the physiological chamber filled with Opti-MEM supplemented with 5% FBS and penicillin-streptomycin (PS). Alternatively, TS were directly placed in the physiological chamber containing medium supplemented with 10 µg/ml of Bleomycin.

### SPIM Setup

The SPIM setup consisted of two perpendicular horizontal optical axes: a light-sheet illumination axis for selective plane excitation and a detection axis. This custom-built SPIM featured a compact laser launch combined with dichroic mirrors into a single multi-wavelength beam. The light sheet was generated using a cylindrical lens conjugated to an illumination objective (10× NA 0.25). The sample was positioned in the light sheet within a chamber filled with an aqueous medium. Light was collected using an immersion objective (20× NA 0.35). The detection objective was housed within a physiological chamber connected to a filter wheel (Lambda 10, Sutter Instrument) and a CMOS camera (Flash 4.0, Hamamatsu). The physiological chamber was created by stereolithography using a photosensitive epoxy resin (Cresilas). Sample movements (x, y, z, and rotation) were managed by a fully motorized stage (Physics Instruments). An incubator controlled the temperature and CO2 concentration around the sample (PECON controllers).

### Sample Holder Preparation for Time-lapse Data Acquisition

Sample holders were made from low melting agarose. Before use, the agarose solution was melted and used to mold the cuvette. The molded cuvette was filled with Opti-MEM (Invitrogen) culture medium containing 5% FBS and 1% penicillin-streptomycin. The TS was placed in the molded cuvette and secured to the SPIM sample holder. This assembly was placed in the physiological chamber filled with Opti-MEM culture medium containing 5% FBS and 1% penicillin-streptomycin.

### Short and Long-term Live Imaging by SPIM

To ensure stability and minimize temperature-induced shifts, the temperature around the SPIM setup was maintained at 37°C for at least 24 hours prior to imaging. For short-term live imaging, TS were imaged over 3 hours with 20-minute intervals. For long-term live imaging, TS were imaged continuously for 50 hours with 1-hour intervals. Imaging parameters included a laser power of 8 mW/cm², an exposure time of 200 ms per image, and Z-stacks of 150 µm, representing a hemispherical section of the TS.

### Imaris

Three-dimensional reconstructions were performed using Imaris 7.0.0 software. The source channel index was set to 1 with estimated diameters of 5.00 µm in XYZ, and background subtraction was enabled. Spots with a “Quality” above 0.600 were filtered for analysis. Region growing was based on absolute intensity with manual thresholding and diameters derived from volume. An intensity StdDev coded colormap spectrum was applied to enhance visualization.

### Quantification and Statistical Analysis

Statistical analysis was performed using Prism 7 (GraphPad Software). Details on data presentation and error bars are provided in the figure legends.

## Data and Resource Availability

### Lead Contact

Further information and requests for resources and reagents should be directed to and will be fulfilled by the lead contact, Corinne Lorenzo.

### Materials Availability

The HCT116 mCherry-mDomain cell line used in this study will be made available upon reasonable request.

### Data and Code Availability

One example of the data sets generated and analyzed during the current study is available on the GitHub repository. All data reported in this paper, including additional raw imaging data, is available from the lead contact upon request. The FIJI macros, 3D texture code, and analysis Jupyter notebook used to generate the results are available in the GitHub repository [https://github.com/corinnelorenzo/4D_DNA_DAMAGE/tree/main]. The computer code for 3D texture was written in C^++^ and must be compiled with Code::Blocks before utilization. A tutorial describing the steps from preprocessing, object identification and classification, to transformation with 3D texture is also available.

Any additional information required to reanalyze the data reported in this paper is available from the lead contact upon request.

## Notes

https://github.com/corinnelorenzo/4D_DNA_DAMAGE/tree/main

### Competing Interest Statement

The authors have declared no competing interest.

